# The Genetic Epidemiology of Developmental Dysplasia of the Hip: A Genome-Wide Association Study Harnessing National Clinical Audit Data

**DOI:** 10.1101/154013

**Authors:** Konstantinos Hatzikotoulas, Andreas Roposch, Karan M Shah, Matthew J Clark, Selina Bratherton, Vasanti Limbani, Julia Steinberg, Eleni Zengini, Kaltuun Warsame, Madhushika Ratnayake, Maria Tselepi, Jeremy Schwartzentruber, John Loughlin, Deborah M Eastwood, Eleftheria Zeggini, J Mark Wilkinson

## Abstract

**Background:** Developmental dysplasia of the hip (DDH) is a common, heritable condition characterised by abnormal formation of the hip joint, but has a poorly understood genetic architecture due to small sample sizes. We apply a novel case-ascertainment approach using national clinical audit (NCA) data to conduct the largest DDH genome-wide association study (GWAS) to date, and replicate our findings in independent cohorts.

**Methods:** We used the English National Joint Registry (NJR) dataset to collect DNA and conducted a GWAS in 770 DDH cases and 3364 controls. We tested the variant most strongly associated with DDH in independent replication cohorts comprising 1129 patients and 4652 controls.

**Results:** The heritable component of DDH attributable to common variants was 55% and distributed similarly across autosomal and the X-chromosomes. Variation within the *GDF5* gene promoter was strongly and reproducibly associated with DDH (rs143384, OR 1.44 [95% CI 1.34-1.56], p=3.55x10^−22^). Two further replicating loci showed suggestive association with DDH near *NFIB* (rs4740554, OR 1.30 [95% CI 1.16-1.45], p=4.44x10^−6^) and *LOXL4* (rs4919218, 1.19 [1.10-1.28] p=4.38x10^−6^). Through gene-based enrichment we identify *GDF5, UQCC1, MMP24, RETSAT* and *PDRG1* association with DDH (p<1.2x10^−7^). Using the UK Biobank and arcOGEN cohorts to generate polygenic risk scores we find that risk alleles for hip osteoarthritis explain <0.5% of the variance in DDH susceptibility.

**Conclusion:** Using the NJR as a proof-of-principle, we describe the genetic architecture of DDH and identify several candidate intervention loci and demonstrate a scalable recruitment strategy for genetic studies that is transferrable to other complex diseases.

**Key Messages:**

- We report the first genome-wide scan for DDH in a European population, and the first to use national clinical audit data for case-ascertainment in complex disease.
- The heritable component of DDH attributable to common variants is 55% and is distributed similarly across autosomal and the X-chromosomes.
- Variation within the *GDF5* gene promoter is strongly and reproducibly associated with DDH, with fine-mapping indicating rs143384 as the likely casual variant.
- Enrichment analyses implicate *GDF5, UQCC1, MMP24, RETSAT* and *PDRG1* as candidate targets for intervention in DDH.
- DDH shares little common genetic aetiology with idiopathic osteoarthritis of the hip, despite sharing variation within the *GDF5* promoter as a common risk factor.

## INTRODUCTION

Developmental dysplasia of the hip (DDH) is characterised by abnormal development of the hip joint and presents with varying severity from mild uncovering of the femoral head to complete dislocation of the joint.^1^ It is the most common developmental musculoskeletal anomaly, with population-weighted average incidence that ranges strongly with ethnic background from 0.06 per 1000 live births in Black Africans to 76.1 per 1000 live births in Native Americans,^2^ and with an incidence in the UK European population of 3.6 per 1000. DDH is a complex disorder, with known associations including female sex, first-born, breech presentation, and family history.^2^ There is a 7-fold increase in the incidence between siblings and a 10-fold increase in the parents of probands compared to the general population,^3^ and a concordance rate of 41% between identical twins versus 3% in dizygotic twins.^4^

Whilst DDH is heritable, its genetic architecture remains poorly characterised. Several linkage scans and candidate gene studies have implicated possible associated genetic variants, including in *GDF5,*^5^ but to date no replicated loci of genome-wide significance have been identified. ^2, 6-9^ A recent genomewide association study (GWAS) of 386 patients and 558 controls in the Han Chinese population suggested an association with variation in *UQCC* (odds ratio (OR) 1.35, *P*=3.63x10^−6^), a gene adjacent to *GDF5*.^10^ Morphological abnormalities of the hip such as DDH are also recognised as risk factors for the development of secondary degenerative change of the hip,^11, 12^ and commonly result in hip replacement in adult life.^12^ ^13^ Idiopathic hip osteoarthritis (OA) is very common and has a substantial heritable component, ^14^ and may share common genetic aetiology with DDH.^2, 7, 15^

Systematic examination of genome-wide variation in DDH in larger sample sizes is necessary to clarify its heritable biology and inform mechanism-driven preventative strategies. However, sample collection for uncommon diseases can be challenging and time-consuming, as attested by the relative paucity of sample sets for genetic studies of DDH. Routinely-collected national clinical audit datasets (NCAs) present an opportunity for efficient case ascertainment in genetic epidemiology. The National Joint Registry for England, Wales, Northern Ireland, and the Isle of Man (NJR, http://www.njrcentre.org.uk/) was established in 2003 to collect audit data on all hip and knee replacement surgery in these regions, for which it has a completeness rate of 97% (http://www.njrreports.org.uk/Data-Completeness-andquality). As at December 31^st^ 2014, the dataset held information on 711 765 primary hip replacement procedures, including data on the diagnostic indication for surgery. We used the NJR as a case-ascertainment tool to conduct a nationwide genome-wide association scan to characterise the genetic architecture of DDH. We examine the heritable contribution to DDH, and use fine-mapping and gene-based enrichment approaches to identify likely causal variants and genes. Finally, we use independent OA-datasets to examine for shared genetic aetiology between DDH and hip OA.

## PATIENTS AND METHODS

### NCA-based recruitment strategy

Between January 2013 and April 2014, 5411 adults (4095 female) from a total NJR database of 711 765 procedures (0.76%) had undergone a hip replacement for the recorded indication of hip dysplasia and were invited to participate in the screening program (figure 1, and supplementary section 1a.). Subsequent screening comprised filtering by self-reported questionnaires for UK European ancestry and a childhood history of idiopathic DDH (supplementary section 1.a and table S1). Following screening, 1091 individuals confirming both UK European ancestry and idiopathic DDH diagnosed in childhood were invited to return by post a saliva sample for DNA extraction and genotyping. Of these, 907 (803 female) returned a sample. The sex distribution of the responding subjects was similar to that of the DDH population in the UK.^2^ Radiographic validation of the DDH phenotype was conducted using a plain pelvic radiograph preceding hip replacement retrieved electronically via the National Data Sharing Network (http://www.image-exchange.co.uk/) in a convenience subset of 231 (25%) subjects completing saliva return. The age and sex distribution of the validation subset was similar to the 907 that returned a saliva sample. Radiographic analysis confirmed DDH in 224/231 (97%, supplementary section 1.a). Non-DDH cases were excluded from further analysis. Samples from 900 subjects proceeded to DNA extraction and quality control. In 834 samples DNA yield was sufficient to progress to genome-wide genotyping.

**Figure 1:**
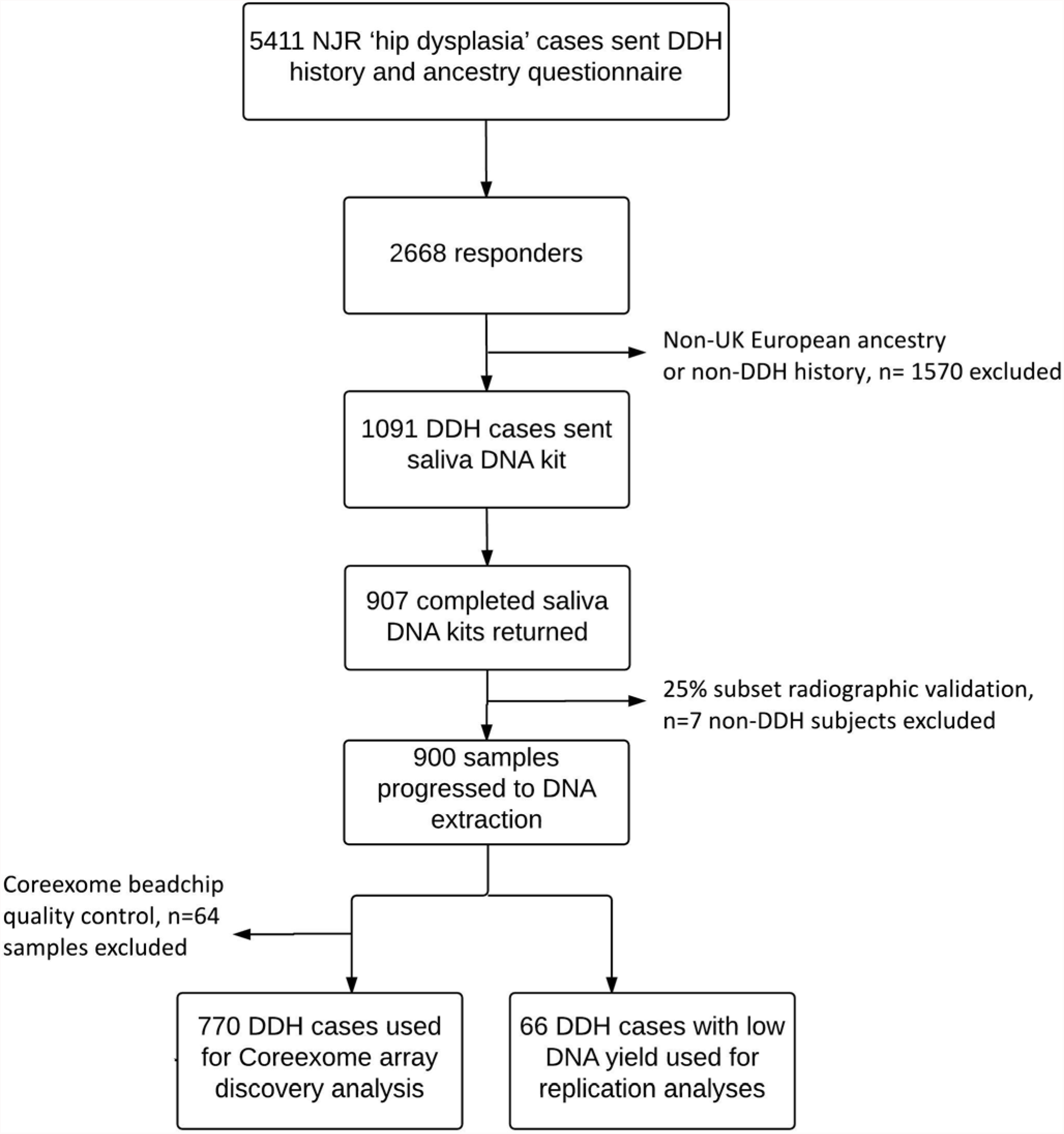
Recruitment flowchart

The replication cohort comprised 838 children (725 female) from the UK DDH case control consortium recruited at 18 English hospitals (supplementary section 1.a and table S2) and 225 adults (220 female) with DDH recruited at the Royal National Orthopaedic Hospital. In addition, 66 cases (59 female) from the NJR discovery cohort of insufficient DNA yield for genome-wide genotyping were also included in the replication cohort. Subject overlap between the discovery and replication populations, and between the replication cohorts was excluded using individual patient identifiers and by cross-referencing to the NJR dataset. Case duplication was excluded in the discovery cohort at both subject ID and genotype quality control stages.

The control population comprised 8016 participants from the United Kingdom Household Longitudinal Study (UKHLS, https://www.understandingsociety.ac.uk/about). A sample of 3364 UKHLS controls was included in the discovery stage and an independent sample of 4652 UKHLS controls was used in the replication stage at a case:control ratio of ~1:4. The proportion of female controls in the discovery stage reflected the distribution in cases. All replication and control participants were of UK European ancestry. The study was approved by the National Research Ethics Service in England (NRES 12/YH/0390, October 30^th^ 2012). All subjects provided written informed consent prior to participation.

### Genotyping, quality control, and association analyses

DNA from 834 NJR DDH cases and UKHLS controls was genotyped using the Illumina HumanCoreExome BeadChip (Illumina, San Diego, USA; supplementary section 1.b and table S3). Power was calculated by fixing the sample size to our post-quality control (QC) sample size and gave >80% power to detect common variants with moderate effect size at genome-wide significance (p=5.0x10^−8^, supplementary section 1.b and figure S1). Following QC checks at both the sample and single nucleotide polymorphism (SNP) level (supplementary section 1.b), 256 867 variants were tested using the likelihood ratio test under an additive genetic model for association with DDH in 770 (693 female) DDH cases and 3364 (3048 female) controls using SNPTEST v2.2.^16^

Independent variants (p<1x10^−4^) (supplementary section 1.b) in the discovery set were selected for replication. *De novo* genotyping for these SNPs in 1129 (1004 female) replication cases was conducted using the iPLEX^®^ Assay and the MassARRAY^®^ System (Agena Bioscience, Inc). Case-control association analysis versus 4652 (2527 female) controls was performed under an additive genetic model. Finally, a fixed-effects meta-analysis was conducted across the discovery and replication datasets, comprising a total of 1899 cases and 8016 controls. Conditional single-variant association analyses were used to identify statistical independence. A variant was considered independent of the index SNP if the pre- and post-conditioning p-value difference was lower than two orders of magnitude.

### Heritability analysis

We estimated the heritability of DDH using the genome-wide analysis dataset using genetic complex trait analysis (GCTA) and phenotype correlation genotype correlation (PCGC), assuming a DDH prevalence of 3.6 per 1000 live births in the UK, ^2^ and using a minor allele frequency (MAF) cut-off of 0.01 (supplementary section 1.c).

### Fine-mapping

In order to fine-map the replicating association signals, we imputed variant genotypes locally using 1000 Genomes Project^17^ combined with UK10K Project^18^ data as the reference panel (supplementary section 1d.). We then applied a method that assigns a relative “probability of regulatory function” (PRF) score among candidate causal variants, reweighting association statistics based on epigenomic annotations, and delineating the 95% probability set of likely causal variants (supplementary section 1.d).

### Gene-based enrichment analyses

In order to quantify the degree of association each gene has with the phenotype and understand the potential functional effects of the identified variants, we used the GWAS results to perform gene-based enrichment analyses in MAGMA v1.03 to identify the joint association of all markers in a given gene with the phenotype (supplementary section 1.e).

### Genetic overlap with hip OA

Polygenic risks scores were created for all genotyped participants to test for shared genetic aetiology with idiopathic hip OA. Two independent UK cohorts were used for these analyses: 3266 cases with hip OA versus 11 009 controls from the arcOGEN study; and 2396 cases with International Classification of Diseases (ICD) 10 coded hip OA versus 9593 controls from the UK Biobank (supplementary section 1.f). High-resolution polygenic risk scores were made using evenly spaced p-value thresholds (Pt) between 0.001 to 0.5, at increments of 0.0001, to produce 4991 risk thresholds.

### Data Availability

Genome-wide genotype data of the DDH cases and UKHLS controls have been deposited to the European Genome-Phenome Archive (www.ebi.ac.uk/ega/home, DDH accession numbers: EGAD00010000766 and EGAS00001000916; UKHLS accession numbers: EGAD00010000891 and EGAS00001001232) and of the DDH NJR cases to the NJR data archive (www.njrcentre.org.uk).

## RESULTS

### The heritable component of DDH

Using GCTA we found that common-frequency autosomal SNPs explain 55% (±se=6%, p<0.0001) of the liability-scale heritability of DDH. The heritability estimate by PCGC was very similar to that identified by GCTA (supplementary table S4). The heritability estimate was also similar (54.7±5.8%) when the analysis was repeated using sex as a covariate.

### Discovery GWAS

Genome-wide analysis showed an excess of signals in the discovery case-control population (figure 2). A total of 53 SNPs comprising 25 independent signals showed suggestive evidence for association with DDH at p<1x10^−4^ compared to 3 independent variants expected under the null hypothesis of no association (binomial p=9.57x10^−17^, supplementary table S5). Eleven correlated variants reached genome-wide significance (p<5.0x10^−8^) and all reside in the same region. The lead variant, rs143384 (effect allele A, effect allele frequency (EAF) 0.60, OR[95%CI] 1.57[1.3-1.77] p=1.72x10^−14^), is located in the 5’ untranslated region of *GDF5* (20q11.22). Since DDH is sex-biased toward females, we also repeated the analysis using gender as a covariate and found no qualitative difference (supplementary table S6).

**Figure 2:**
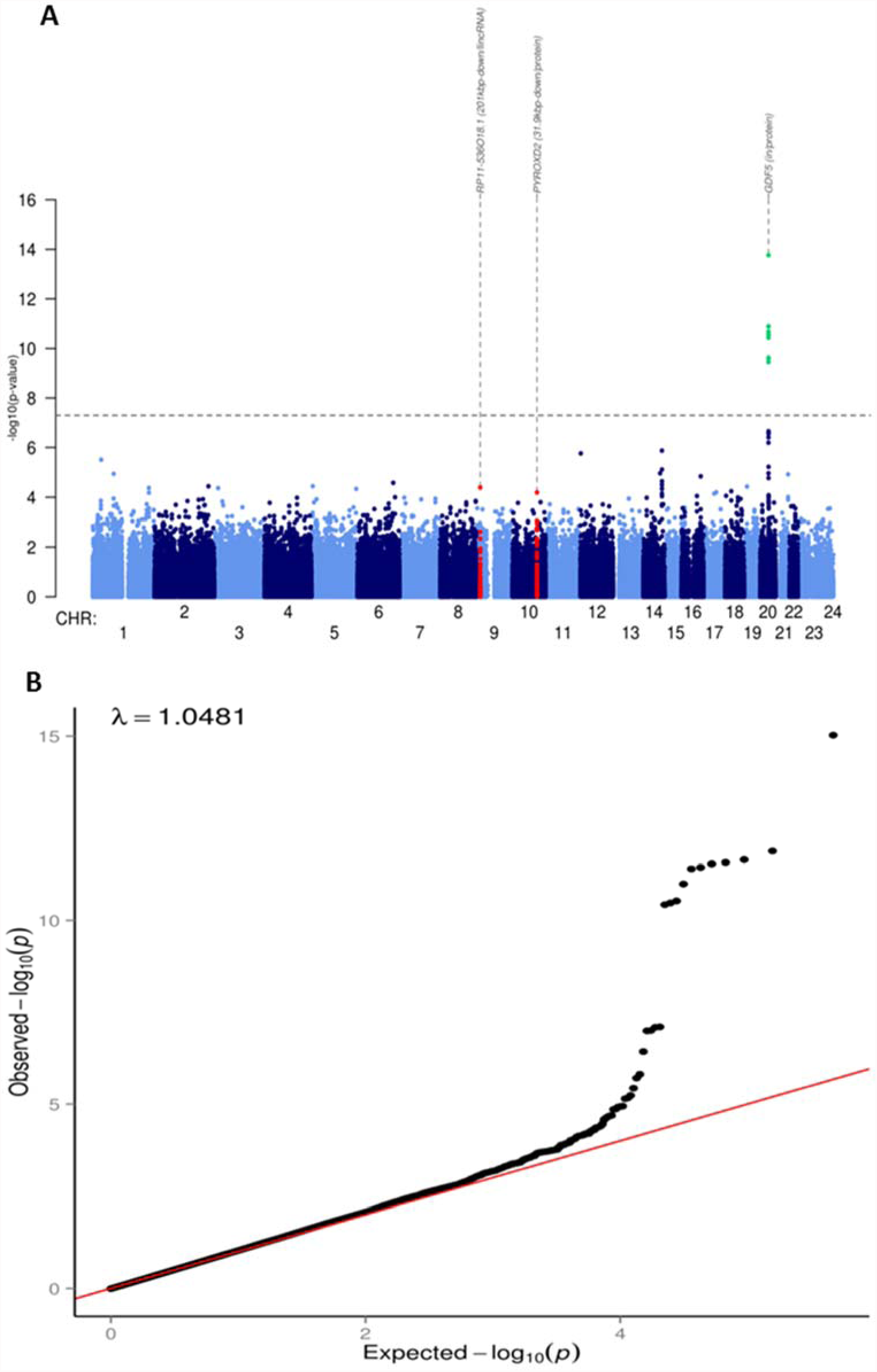
Manhattan and quantile-quantile plot of the DDH genome-wide association scan. Panel A is a Manhattan plot showing the -log_10_ p values for each variant (y axis) plotted against their respective chromosomal position (x axis). The dashed line indicates the genome-wide significance threshold (p=5.0×10^−8^). Green dots represent variants for which p values reached the genome-wide significance threshold where red dots illustrate the other two replicating signals. Panel B is a quantile-quantile plot. The x-axis indicates the expected −log_10_ p values and the y-axis the observed ones. The red line represents the null hypothesis of no association at any locus.

### Replication

Replication of the independent signals was conducted in three further DDH populations of UK European ancestry, totalling 1129 cases and 4652 UKHLS controls. The variant rs143384 reached genome-wide significance in the replication dataset (effect allele A, EAF 0.61, OR 1.37[1.24 to 1.51] p=1.33x10^−10^). Four further SNPs showed nominal association (p<0.05) with the same direction of effect as the discovery cohort, compared with 1.35 under the null expectation (binomial test p=0.01, supplementary table S7). When the replication analysis was performed excluding the 66 NJR subjects that failed genome-wide genotyping QC, the findings were similar (data not shown).

### Meta-analysis

At meta-analysis, 20 SNPs were associated with DDH with the same direction of effect in both the discovery and replication analysis. Three SNPs in the same region of chromosome 20 showed association with DDH at genome-wide significance: rs143384 in *GDF5* (figure 3A, effect allele A, OR 1.44[1.34 to 1.56] p=3.55x10^−22^), rs12479765 in *MMP24* (effect allele G, OR 1.33[1.20 to 1.47] p=3.18x10^−08^), and rs2050729 in *RMB29* (effect allele G, OR 1.41[1.25 to 1.58] p=1.15x10^−08^). Conditional analysis of rs12479765 and rs2050729 on the lead variant rs143384 attenuated their association with DDH, indicating that they were conditionally dependent upon the lead variant. rs143384 explained 0.96% of DDH variance on the liability scale (h^2^L) and the sibling relative risk ratio (*λ*_s_) attributable to this variant was *λ*_s_=1.04 (supplementary figure S2).

**Figure 3:**
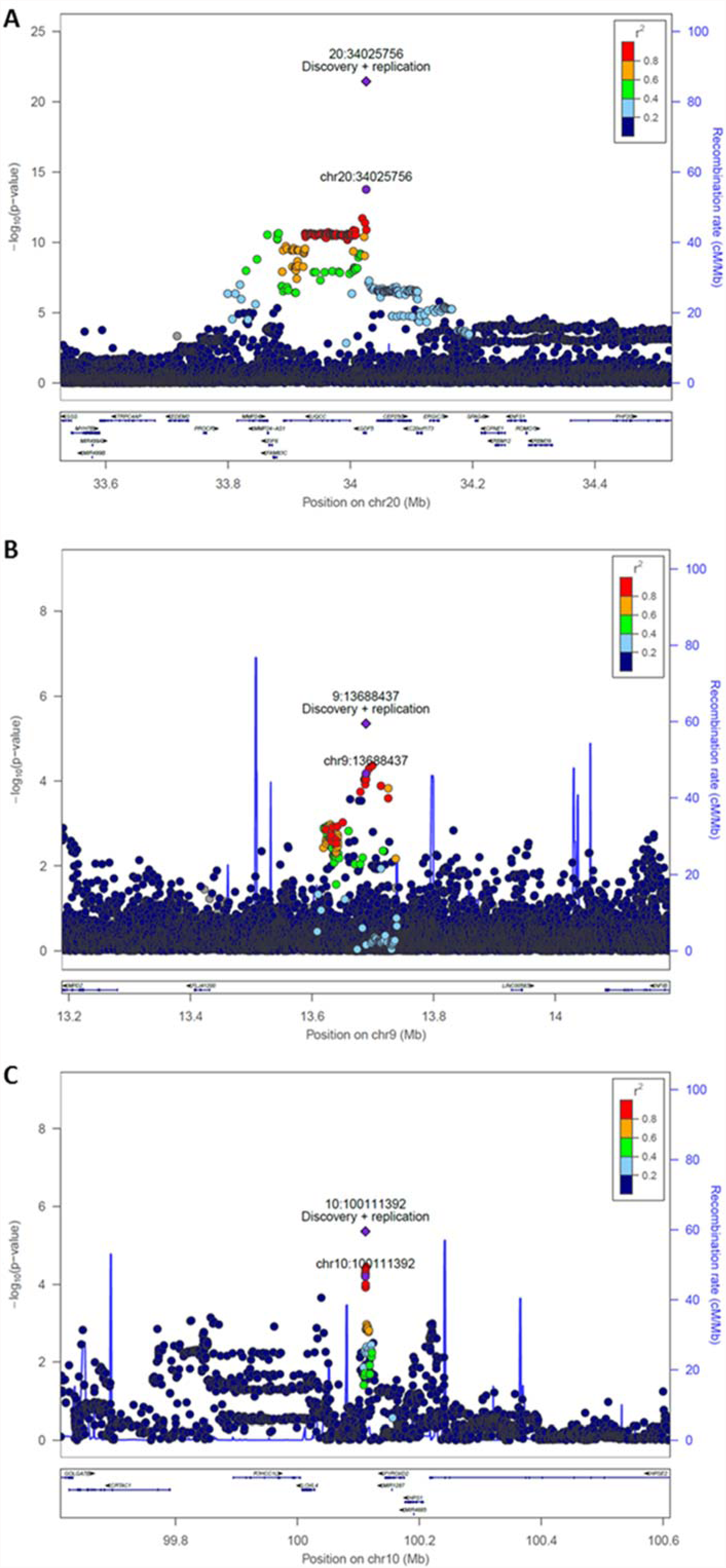
Region association plots of the three replicating signals. Regional association plot of rs143384 (20:34025756, panel A), rs4740554 (9:13688437, panel B) and rs4919218 (10:100111392, panel C). Each filled circle represents the p value of analysed variants (as -log_10_ values) plotted against their physical position (NCBI Build 37). The p value at the discovery stage and combined discovery and replication cohorts is represented by a purple circle and diamond, respectively. The rest variants in the region are coloured depending on their degree of correlation (r^2^) with the index variant according to a scale from r^2^=0(blue) to r^2^=1(red). Underneath, gene annotations from the UCSC genome browser.

Two further independent signals showed evidence of replicating association with DDH, although this did not reach genome-wide significance at meta-analysis: rs4740554 (effect allele C, EAF 0.10, OR 1.30[1.16 to 1.45] p=4.44x10^−6^) and rs4919218 (effect allele C, EAF 0.31, OR 1.19[1.10 to 1.28] p=4.38x10^−6^) (figure 3B and 3C, and Table 1). rs4740554 and rs4919218 explained 0.36% and 0.24% of phenotypic variance (*λ*_s_= 1.02 and 1.01), respectively (supplementary figure S2).

**Table 1:** Replicating variants associated with DDH.

| **SNP ID**<br>**(chromosome:position)** | **Effect**<br>**Allele/Other**<br>**Allele** | **Effect**<br>**Allele**<br>**Frequency**<br>**(discovery**<br>**stage)** | **Discovery**<br>**Odds Ratio**<br>**(95% CI)** | **Discovery**<br>**p value** | **Replication**<br>**Odds Ratio**<br>**(95% CI)** | **Replication**<br>**p value** | **Meta-**<br>**analysis**<br>**Odds Ratio**<br>**(95% CI)** | **Meta-**<br>**analysis**<br>**p value** |
| --- | --- | --- | --- | --- | --- | --- | --- | --- |
| rs143384 (20:34025756) | A/G | 0.60 | 1.57<br>(1.4-1.77) | 1.72E-14 | 1.37<br>(1.24-1.51) | 1.33E-10 | 1.44<br>(1.34-1.56) | 3.55E-22 |
| rs4740554 (9:13688437) | C/A | 0.10 | 1.44<br>(1.22-1.71) | 4.09E-05 | 1.20<br>(1.03-1.39) | 0.01 | 1.30<br>(1.16-1.45) | 4.44E-6 |
| rs4919218<br>(10:100111392) | C/T | 0.69 | 1.27<br>(1.13-1.43) | 6.48E-05 | 1.14<br>(1.03-1.25) | 0.0089 | 1.19<br>(1.10-1.28) | 4.38E-6 |

### Fine-mapping

For the *GDF5* signal, five variants were included in the 95% probability set of being causal. Among these, both rs143384 and rs143383 have high epigenomic scores as they are located in the 5’ UTR of *GDF5*, have high nucleotide sequence conservation, and overlap DNase hypersensitivity and activating histone marks. Considering annotations and association statistics p-values together, rs143384 had a >99% likelihood of being the causal variant, assuming only 1 variant as causal.

### Gene-based enrichment analyses

After adjusting for the family-wise error rate (FWER), we found five genes that were associated with DDH susceptibility: *GDF5*, *UQCC1*, *MMP24*, *RETSAT* and *PDRG1* (p=9.24x10^−12^, p=1.86x10^−10^, p=3.18x10^−9^, p=3.70x10^−8^ and p=1.06x10^−7^, respectively; Supplementary table S8). To exclude association through *GDF5*, we repeated the analysis by removing *GDF5* and all genes remained associated with DDH.

### Shared genetic aetiology with hip OA

We found little evidence for shared genetic aetiology between DDH and hip OA. The best fit polygenic risk score for hip OA in the arcOGEN and ICD-10 coded hip OA in the UK Biobank datasets explained ~0.05% and 0.35%, respectively, of the variation in DDH susceptibility (FWER P>0.05, supplementary figure S3).

## DISCUSSION

We used case-ascertainment by NCA database search and postal recruitment to facilitate the largest genome-wide association scan for DDH to date. We found that the heritable component of DDH due to common autosomal variants is approximately 55%, consistent with the complex nature of the disease. We establish variation within *GDF5* on chromosome 20 as robustly associated with DDH susceptibility, with the variant rs143384 as the causal signal by fine-mapping, although *GDF5* variation makes only a small contribution to overall DDH heritability. Through gene-based enrichment analyses we identify *GDF5, UQCC1, MMP24, RETSAT* and *PDRG1*, as independently associated with DDH susceptibility.

Previous studies examining the epidemiology of DDH within families using epidemiological linkage approaches have demonstrated that heritable factors contribute between 50% and 85% of the total liability of disease,^3, 19, 20^ depending on population ethnicity. Here, we used dense genome-wide genotyping data to estimate the heritable risk of DDH amongst unrelated individuals of UK European ancestry in the general English population. Our estimate of DDH heritability attributable to variation within autosomal chromosomes was broadly similar to the estimates derived from linkage analyses.^2^ When the heritability analysis was stratified by chromosome, we found that the X-chromosome contributed a similar amount to overall DDH heritability as the autosomes (Supplementary Fig. 4), and we identified no individual chromosome X signals. Taken together, these data suggest that although DDH is strongly sex-linked, this is not substantially due to variation in genomic DNA.

The gene *GDF5* encodes growth differentiation factor 5 (GDF5), belonging to the transforming growth factor beta superfamily.^21^ GDF5 is required for normal bone and joint development by promoting cartilage condensation and increasing the size of the skeletal elements through proliferation within epiphyseal cartilage.^22^ Two *GDF5* variants, rs143383 and 143384, have previously shown suggestive association with DDH in unreplicated candidate gene studies in Asians and in Europeans. ^5, 23^ Here we validate and extend these findings at the genome-wide level and with robust replication. These variants are in high linkage disequilibrium (r^2^=0.82, D’=0.99), and were both directly genotyped in this study. Our data implicate rs143384 as the lead variant, with the rs143383 association with DDH being two orders of magnitude weaker. Subsequent fine-mapping of the *GDF5* locus using Roadmap Epigenome annotations with imputation using 1000 Genomes and UK10K Project data also identified rs143384 as the likely causal variant. Chen et al,^24^ have shown that the *GDF5* locus contains many separate regulatory elements that control expression of the gene at different joint sites, and that these flanking regions are large. Interrogation of the GTEx database indicates that rs143384 is an expression quantitative trait locus (eQTLs) for multiple genes across various tissues (supplementary table 9).^25, 26^ This annotated function provides novel opportunities for investigation of the role of GDF5 as a candidate target for DDH prevention.

Gene-based enrichment analysis can boost power to identify contributing loci by combining information across multiple SNPs co-localised at the gene level. Here, we found associations with the *GDF5, UQCC1, MMP24, RETSAT* and *PDRG1* genes. *UQCC1* gene encodes a trans-membrane protein ubiquinol-cytochrome c reductase complex chaperone, which is structurally similar to the mouse basic fibroblast growth factor repressed ZIC-binding protein. ^27^ In humans, polymorphisms in this gene have been associated with DDH,^10^ bone size,^28^ height,^29^ and hip axis length.^30^ *MMP24* encodes a member of the peptidase M10 family of matrix metalloproteinases (MMPs) involved in the breakdown of extracellular matrix in normal physiological processes, such as embryonic development and tissue remodelling.^31^ Polymorphisms within *MMP24* have also been associated with height variation in childhood. ^32, 33^ *RETSAT* codes for retinol saturase, an enzyme centrally involved in the metabolism of vitamin A.^34^ Retinoic acid signalling is essential for normal limb bud development, including bone and cartilage formation, ^35^ and provides a further candidate target in DDH pathogenesis.

This study has limitations. The patients recruited in the discovery cohort were adults who had undergone a hip replacement for progressive symptoms related to an underlying diagnosis of hip dysplasia, as identified through the NJR. To mitigate against misclassification bias in case-ascertainment we employed a stepwise filtration approach that subsequently validated with high concordance against examination of plain radiographic images. The female-to-male case ratio in our final NJR DDH cohort was similar to that expected in the general DDH population in the UK,^2^ and contrasts with that found in idiopathic OA of the hip.^36^ Previous studies have also demonstrated the contribution of DDH as a risk factor for secondary OA,^11, 12, 37^ and that OA susceptibility genes influence the association between hip morphology and OA. ^15, 38, 39^ We therefore examined whether we were simply detecting OA-associated loci in our DDH discovery cohort, as case identification required NJR registration for a hip replacement. Polygenic risk scores derived using independent hip OA cohorts predicted only a small amount of the genetic variability in the DDH discovery population, suggesting only limited common genetic aetiology between the diseases. Finally, our replication cohorts mainly comprised independent populations of children recruited prospectively with a firm diagnosis of DDH, and in whom the odds-ratio of association for the primary variant signal was similar to that found in the NJR-discovery sample.

In summary, the results of this largest study of DDH to date provide a comprehensive picture of its complex genetic architecture, the contribution of common variants to its heritability, and establishes the first robust DDH genetic locus. We also demonstrate the utility of national clinical audit-based case ascertainment and recruitment for genetics and genomics studies using a strategy that is transferrable to other complex diseases in which large-scale datasets are held.

## Acknowledgments

We thank the patients and staff of all the hospitals who have contributed data to the National Joint Registry. We are grateful to the Healthcare Quality Improvement Partnership (HQIP), the National Joint Registry Steering Committee (NJRSC), and staff at the NJR Centre for facilitating this work. The views expressed represent those of the authors and do not necessarily reflect those of the NJRSC or HQIP who do not vouch for how the information is presented. This research has been conducted using the UK Biobank Resource under Application Number 9979. The study also utilised summary genotype data from arcOGEN (http://www.arcogen.org.uk/) funded by a special purpose grant from Arthritis Research UK (grant 18030).

